# Feasibility of Administering Human Pancreatic Cancer Chemotherapy in a Spontaneous Pancreatic Cancer Mouse Model

**DOI:** 10.1101/2021.09.08.459476

**Authors:** Abagail M. Delahoussaye, Joseph Abi Jaoude, Tara N. Fujimoto, Jessica Molkentine, Carolina J. Garcia Garcia, Jason P. Gay, Ningping Feng, Joe Marszalek, Cullen M. Taniguchi

## Abstract

**Background:** Both modified FOLFIRINOX (mFFX) and gemcitabine/*nab*-paclitaxel chemotherapy regimens have been shown to improve clinical outcomes in patients with pancreatic cancer, and are often used interchangeably as the standard of care. Preclinical studies often do not use these regimens, since administering these multiagent approaches can be difficult. In this study, we assessed the feasibility of administering these two chemotherapy regimens in spontaneous pancreatic tumors using KPC mice with the ultimate goal of advancing preclinical studies.

**Methods:** KPC mice were created by breeding *Kras*^*LSL-G12D/+*^ to *Trp53*^*fl/fl*^;*Ptf1α*^*Cre/+*^, resulting in *Kras*^*LSL-G12D/+*^;*p53*^*fl/+*^;*Ptf1α*^*Cre/+*^ mice. At 14 weeks of age, mice were palpated for spontaneous tumor growth that was verified using ultrasounds. Mice with tumors under 15 mm in diameter were used. The mice were assigned to one of seven treatment regimens: 1 cycle of mFOLFIRINOX (FFX X1), 2 cycles of mFOLFIRINOX (FFX X2), 1 cycle of mFOLFIRINOX with 40Gy SBRT (FFX SBRT), 1 cycle of gemcitabine/*nab*-paclitaxel (GEM/AB X1), 2 cycles of gemcitabine/*nab*-paclitaxel (GEM/AB X2), 2 cycles of gemcitabine/*nab*-paclitaxel with 40Gy SBRT (GEM/AB SBRT), or no chemotherapy treatment (control).

**Results:** In total, 92 mice were included. The median OS in the FFX X2 group was slightly longer that the median OS in the FFX X1 group (15 days vs 11 days, *P*=0.003). Mice in the GEM/AB X2 group had longer OS when compared to mice in the GEM/AB X1 group (33.5 vs 13 days, *P*=0.001). Mice treated with chemotherapy survived longer than untreated control animals (median OS: 6.5 days, *P* <0.001). Moreover, in mice treated with chemotherapy, mice that received 2 cycles of GEM/AB X2 had the longest survival, while the FFX X1 group had the poorest OS (*P* <0.001). Lastly, chemotherapy followed by consolidative SBRT trended towards increased local control and survival.

**Conclusions:** We demonstrate the utility and feasibility of clinicall relevant mFOLFIRINOX and gemcitabine/*nab*-paclitaxel in preclinical models of pancreatic cancer.

## Background

Pancreatic cancer is one of the leading causes of cancer-related death in the world [1, 2]. Owing to the limited role of screening in pancreatic cancer, tumors are often discovered at advanced stages, which contributes to poor outcomes and severe morbidity and mortality. While surgery is the main curative option in pancreatic cancer, a large proportion of patients with pancreatic cancer are initially diagnosed with unresectable disease that would not benefit from local therapy [1, 3]. Moreover, pancreatic cancer is known to metastasize early in the disease process [4]. Therefore, chemotherapy remains a major part of treatment for all stages of pancreatic cancer.

Many chemotherapy regimens have been suggested for the treatment of pancreatic cancer. Previously, gemcitabine and 5-fluorouracil were used as monotherapy, however recurrence rates remained high [5, 6]. As such, modern systemic treatment consists \of powerful combinations of chemotherapy. The two most common regimens used in the modern treatment of pancreatic cancer are modified FOLFIRINOX (mFFX: oxaliplatin, irinotecan, leucovorin, fluorouracil) and gemcitabine/*nab*-paclitaxel [7–10]. Many studies and trials analyzing the role of those regimens showed superior clinical results compared to conventional monotherapy treatments, and established the role of mFOLFIRINOX and gemcitabine/*nab*-paclitaxel as standard of care in pancreatic cancer.

While those regimens have been thoroughly studied in humans, limited data on the use of mFOLFIRINOX and gemcitabine/*nab*-paclitaxel are present in pre-clinical studies. In particular, the clinical utility and toxicity of those regimens is still not clear in spontaneous pancreatic tumors in KPC mice. In order to properly mimic human clinical treatment, pre-clinical models need to implement similar chemotherapy regimens. In this study, we aim to assess the feasibility of administering mFOLFIRINOX and gemcitabine/*nab*-paclitaxel in spontaneous pancreatic tumors using a KPC mouse model, as well as their toxicity profiles.

## Methods

### KPC mice and tumor diagnosis

All mouse work was approved and done in accordance with the Institutional Animal Care and Use Committee of The University of Texas MD Anderson Cancer Center under protocol IACUC #00001252-RN02. The studies were carried out in compliance with ARRIVE guidelines. Both female and male mice were used in this study. Mice were maintained on a 12-hour light/dark cycle and were provided with sterilized water and standard rodent chow (Prolab Isopro RMH 3000 irradiated feed). Experiments were carried out during the light cycle.

KPC mice were created by breeding *Kras*^*LSL-G12D/+*^ to *Trp53*^*fl/fl*^;*Ptf1α*^*Cre/+*^, resulting in a *Kras*^*LSL-G12D/+*^;*Trp53*^*fl/+*^;*Ptf1α*^*Cre*^ (KPC) mice, which is a validated model of locally advanced pancreatic cancer [11]. Beginning at 14 weeks of age, mice were screened weekly by palpation under anesthesia for tumor growth, as previously described [12]. Suspicious intrabdominal masses, were then subjected to small animal ultrasound imaging (FujiFilm Vevo2100), with a 30MHz transducer, acquired in B-MODE to estimate volumes from the long and short axes [12, 13].

### Chemotherapy

We used pharmaceutical grade mFOLFIRINOX that was provided by The University of Texas MD Anderson Cancer Center pharmacy. Oxaliplatin was administered intravenously via the tail vein at 10 mg/kg in a volume of 0.01 mL/g. 5-FU (Fresenius Kabi, 50 mg/mL) and Irinotecan (TEVA, 20 mg/mL) were combined and diluted with 0.9% Sodium Chloride (normal saline) and given via intraperitoneal injection at 50 mg/kg. Both injections account for one cycle delivered in the same day. For mice that received two cycles of mFOLFIRINOX, the second cycle was given 2 weeks after completion of the first cycle (Supplementary Figure 1).

For mice treated with gemcitabine/nab-paclitaxel, we used pharmaceutical grade gemcitabine (Gemzar®, NDC 00002-7501) and *nab*-paclitaxel (Abraxane®, Celgene) generously provided by The University of Texas MD Anderson Cancer Center Institute for Applied Cancer Science (IACS). Gemcitabine was diluted with saline to 10 mg/mL and administered with intraperitoneal injections at 100 mg/kg in a volume of 0.01 mL/g, and given twice weekly. *Nab*-paclitaxel (Abraxane®, Celgene) was diluted with saline to 30 mg/mL and administered intravenously at a dose of 300 mg/kg (30mgPTX/kg) in a volume of 0.01 mL/g, and given only once weekly one hour after one of the gemcitabine doses [14] (Supplementary Figure 1).

### Chemotherapy dose finding and efficacy

Mice with detectable tumors were randomly assigned to receive modified FOLFIRINOX (mFFX) or gemcitabine/*nab*-paclitaxel (GEM/AB), and mice received therapy within 48 hours of assignment. Both mFFX and GEM/AB arms were initially assigned to received one cycle of chemotherapy (FFXx1 and GEM/AB x1). If no excessive toxicity was observed, then the next batch of mice would receive 2 cycles of chemotherapy (FFXx1 and GEM/AB x1). No further treatment was given after the last assigned dose of chemotherapy. Body weights were measured daily, and tumor growth monitored twice weekly with ultrasound. Endpoints were reached when mice met predetermined euthanasia endpoints, including tumors measuring greater than 15 mm in diameter, lethargy, weight loss exceeding 20% from baseline, or appearing moribund. At euthanasia, necropsies were conducted to look for macrometastases and determine a potential cause of death, as we have previously described [13].

### Radiation Therapy

For mice that received local treatment with radiation therapy, stereotactic body radiation therapy (SBRT) was delivered using an XRAD 225Cx irradiator with isoflurane anesthesia manifold and image guidance. The tumors were palpated and their locations marked with metallic beads. Then, radiation fields were aligned for each mouse using cone beam CT image guidance. The metallic beads were removed before radiation. Beam arrangement was anteroposterior/posteroanterior using a 10 mm diameter collimator, and mice received 5 fractions of 8Gy with 24 hours between fractions, for a total of 40Gy [13]. mFOLFIRINOX SBRT mice were irradiated two weeks following one dose of chemotherapy. Gemcitabine/*nab-*paclitaxel SBRT mice were given two doses of chemotherapy and then received SBRT two weeks after the final dose. Mice were allowed to feed and drink water *ad libitum* throughout the duration of treatments. Additional supportive measures, such as moistened food pellets on the ground, were given if the mouse appeared dehydrated. Mice were also observed for radiation toxicity daily. The main symptoms assessed were the development of ruffled fur, persistent diarrhea, and/or drastic weight loss of 20% or more.

### Statistical Analysis

Descriptive statistics were used to assess the baseline characteristics of mice and tumors in each group of mice. Continuous variables were presented as median and corresponding interquartile range (IQR). Categorical variables were presented as frequencies and percentages. We further assessed the average weight of mice in each group every day post-treatment. Furthermore, survival curves for Overall Survival (OS) were generated using the Kaplan-Meier method. OS was calculated from time of diagnosis until death of mice. Local tumor recurrence, distant metastasis, and cause of death were noted for all mice. Statistical significance was set *a priori* at *P* <0.05. All statistical analyses were performed using IBM SPSS version 26. Kaplan-Meier curves were generated using Prism Version 8.

## Results

### Baseline Mice and Tumor Characteristics

In total, 92 mice were included and analyzed in our study. In particular, 54 mice were treated with chemotherapy only: 6 mice were treated with one cycle of mFOLFIRINOX (FFX X1), 30 mice with two cycles of mFOLFIRINOX (FFX X2), 6 mice with one cycle of gemcitabine/*nab*-paclitaxel (GEM/AB X1), 6 mice with two cycles of gemcitabine/*nab*-paclitaxel (GEM/AB X2), and 28 mice did not receive any chemotherapy (control).

Table 1 presents the baseline characteristics of mice and tumors in mice treated with chemotherapy only. Mice in the FFX X2 group had the oldest age at diagnosis (24.8 weeks, IQR [21.7-26.9]), and those in the GEM/AB X2 had the youngest age at diagnosis (14.3 weeks, IQR [9.1-23.5]). A similar proportion of males and females were present in the FFX groups, and a few more males than females were present in the GEM/AB and control groups. Baseline weights were similar between all cohorts (between median: 23.6g, IQR 20.5-28.1 and median: 25.3g, IQR 22.0-28.4).

The baseline tumor diameter was largest in the FFX X1 group (median: 9.3 mmm IQR 7.1-10.2), smallest in the GEM/AB X2 group (median: 4.8 mm, IQR 4.0-6.9), and similar in the other 3 groups (between median: 6.7 mm, IQR 6.2-7.4 and median: 7.1 mm, IQR 5.5-8.0). Most tumors were located in the body of the pancreas for the FFX X1 group (3/6, 50.0%), the tail of the pancreas for the FFX X2 group (13/30, 43.3%), the head of the pancreas for the GEM/AB X1 group (4/6, 66.7%), the tail of the pancreas for the GEM/AB X2 group (3/6, 50.0%), and the head of the pancreas for the control group (13/28, 46.4%) (Table 1).

### Comparative analysis between mFOLFIRINOX and gemcitabine/nab-paclitaxel reveals similar survival times

We ultimately wanted to treat KPC animals with at least two cycles of chemotherapy, since that might better approximate the multicycle regimens given to patients [7]. Thirty-six mice were treated with mFOLFIRINOX (6 mice with one cycle and 30 mice with two cycles). The median OS in the FFX X2 group was slightly longer that the median OS in the FFX X1 group (15 days vs 11 days, log-rank *P*=0.003) (Fig. 1A). Twelve mice were treated with gemcitabine/*nab*-paclitaxel (6 mice with one cycle and 6 mice with two cycles). Mice in the GEM/AB X2 group had longer OS when compared to mice in the GEM/AB X1 group (median OS, GEM/AB X2: 33.5 days vs GEM/AB X1: 13 days, log-rank *P*=0.001) (Fig. 1B).

**Figure 1.**
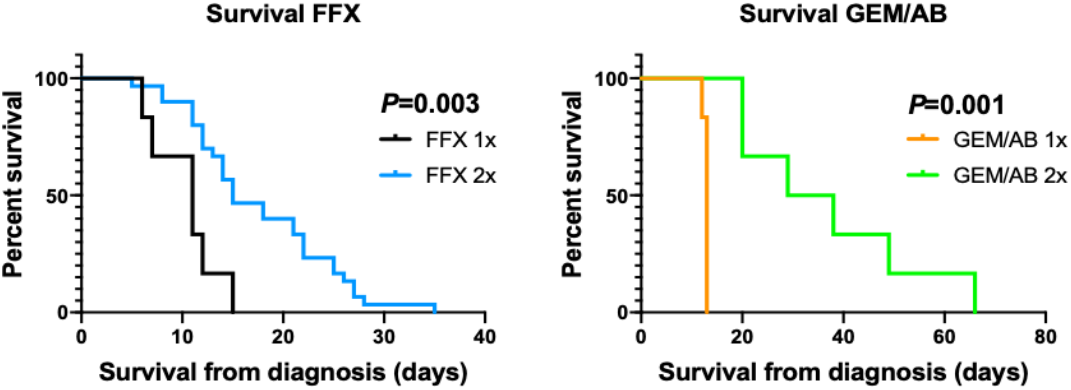
Kaplan-Meier Curve for Overall Survival in Mice Treated with One or Two Cycles of mFOLFIRINOX (FFX) (A), and in Mice Treated with One or Two Cycles of gemcitabine/*nab*-paclitaxel (GEM/AB) (B).

We compared OS between the mFFX and the GEM/AB treated mice. In mice that received one cycle of chemotherapy, the GEM/AB group had a slightly longer median OS compared to the FFX group, but this difference was not statistically significant (GEM/AB X1: 13 days, FFX X1: 11 days, log-rank P=0.104) (Fig. 2A). In mice that received two cycles of chemotherapy, the GEM/AB mice had better OS than the FFX mice (median OS, GEM/AB X2: 33.5 days, FFX X2: 15 days, log-rank *P*=0.002) (Fig. 2B). All mice treated with chemotherapy showed a better median OS compared to the control group (median OS: 10.0 days, log-rank *P* <0.001) (Fig. 2C). Overall, in mice treated with chemotherapy, the GEM/AB X2 group showed the best OS, and the FFX X1 group had the poorest results (log-rank *P* <0.001) (Fig. 2C).

**Figure 2.**
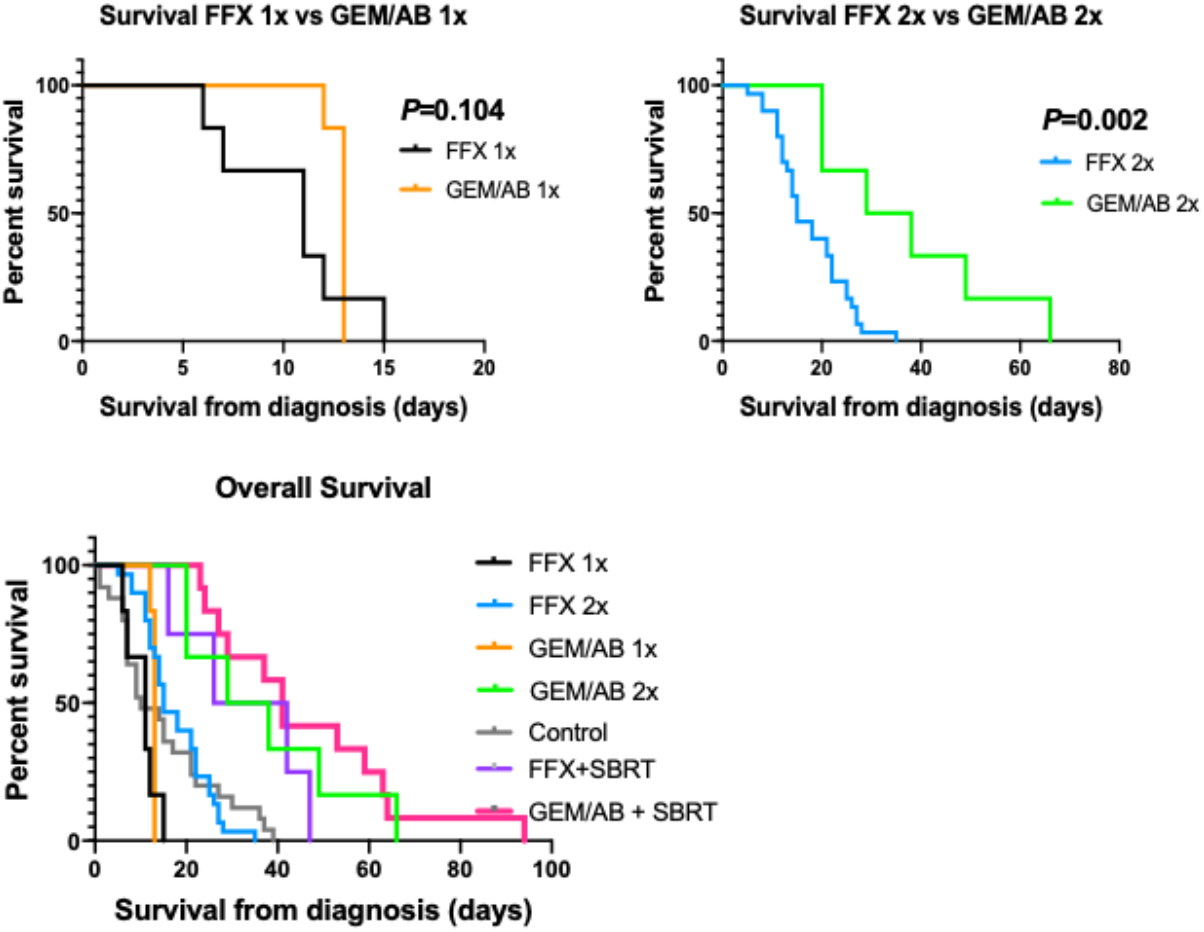
Kaplan-Meier Curve for Overall Survival in Mice Treated with One Cycle of mFOLFIRINOX (FFX) or gemcitabine/*nab*-paclitaxel (GEM/AB) (A), and in Mice Treated with Two Cycles of mFOLFIRINOX (FFX) or gemcitabine/*nab*-paclitaxel (GEM/AB) (B), and Among All Mice (C).

Our KPC animals are a validated model of autochthonous and localized pancreatic cancer. Thus, not surprisingly, local progression was the main cause of death in both FFX groups (FFX X1: 5, 83.3%; FFX X2: 20, 66.7%, Table 2), compared to distant metastasis (FFX X1: 1, 16.7%; FFX X2: 11, 36.7%, Table 2). A similar pattern emerged in GEM/AB treated mice, where local progression of disease led to death more commonly than distant metastases (Table 2). Particularly, in the GEM/AB X1 cohort, 4 (66.7%) mice died from local recurrence, and 1 (16.7%) mouse died from distant metastasis. Moreover, in the GEM/AB X2 cohort 4 (66.7%) mice died from local recurrence, and 2 (33.3%) died from distant metastasis. In fact, ascites was observed in 7 (7/36, 14.4%) of mFFX and 1 (1/12, 8.3%) of GEM/AB mice.

We noted no major differences in acute toxicity between the treatment regimens, as assessed by weight changes (Fig. 3). The average weight of mice in the GEM/AB X1 and FFX X1 decreased over time, while that of the GEM/AB X2 and FFX X2 increased.

**Figure 3.**
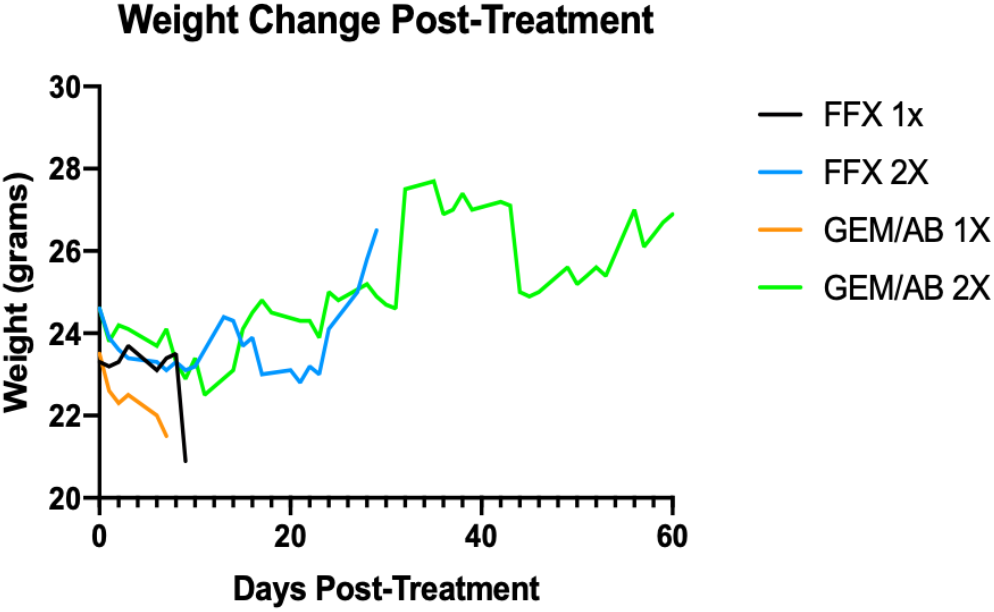
Mice Weight Change After Chemotherapy Treatment.

### Chemoradiation may be superior to chemotherapy alone in KPC mouse models

Radiation therapy is a commonly prescribed to consolidate treatment of the primary mass in pancreatic cancer, especially if surgery cannot be offered [15]. Stereotactic body radiation therapy (SBRT) has emerged as a convenient and powerful methods to achieve local control [16]. We previously demonstrated that SBRT is feasible in murine models [17]. To this end, we treated cohorts of KPC mice that first received mFOLFIRINOX or gemcitabine/*nab*-paclitaxel with a clinically relevant dose of SBRT (8Gy x 5 fx). Sixteen mice were treated with chemoradiation: 4 mice (25%) were treated with one cycle of mFOLFIRINOX and SBRT, and 12 mice (75%) were treated with two cycles of gemcitabine/*nab*-paclitaxel and SBRT.

Table 3 presents the baseline characteristics of mice treated with consolidative SBRT after induction chemotherapy. The median OS in the FFX + SBRT group was 34 days, and the median OS in the GEM/AB + SBRT group was 41 days (log-rank *P*=0.214) (Fig. 4). The addition of SBRT to both mFFX and GEM/AB regimens seems to show a trend towards improved survival compared to mice treated with chemotherapy only (Fig. 2). In total, 4 (25%) mice treated with chemoradiation died from local recurrence: 3 in the FFX group (3/4, 75%), and 1 in the GEM/AB group (1/12, 8.3%) (Table 4). Only one mouse (8.3%) in the GEM/AB group died from radiation toxicity.

**Figure 4.**
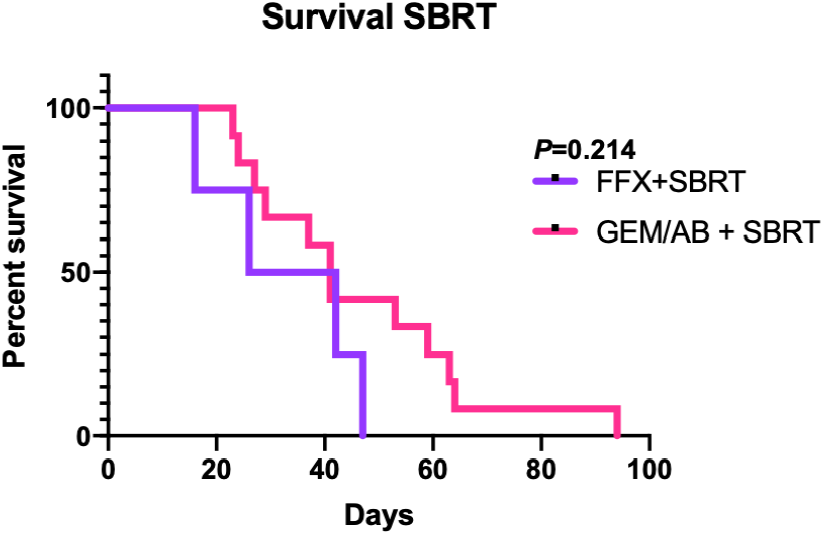
Kaplan-Meier Curve for Overall Survival in Mice Treated Stereotactic Body Radiation Therapy (SBRT) and mFOLFIRINOX (FFX) or gemcitabine/*nab*-paclitaxel (GEM/AB).

## Discussion

The use of chemotherapy is common in patients with pancreatic cancer since most tumors present with advanced or metastatic disease where local therapy is of limited clinical benefit. Two common chemotherapy regimens in the modern treatment of pancreatic cancer are mFOLFIRINOX and gemcitabine/*nab*-paclitaxel [8, 9, 18, 19]. However, the use of those regimens is not routinely performed in pre-clinical models that often inform targets for clinical trials. In our study, we show that treatment with mFOLFIRINOX and gemcitabine/*nab*-paclitaxel is feasible and prolong survival. Thus, the routine use of clinical relevant chemotherapy should be considered in studies involving spontaneous pancreatic tumors in KPC mice.

When analyzing mice that were treated with chemotherapy only, we found that mice treated with two cycles of gemcitabine/*nab*-paclitaxel had the best survival outcomes (median OS: 33.5 days, Fig. 2C). A major component of this regimen is the nanoparticle-albumin-bound paclitaxel which contains albumin, a human derived protein [20]. As such, using *nab*/paclitaxel in KPC mice could be postulated to lead to significant toxicity due to cross reactivity with the human component of the drug with repeated doses. However, we show that gemcitabine/*nab*-paclitaxel was very well tolerated by KPC mice even with repeated dosing, and even led to improved OS compared to mFOLFIRINOX (Fig. 2A, 2B). This similar effect was observed in prior studies, but it is unclear whether prolonged administration of GEM/AB could be feasible. When assessing weight change, mice treated with gemcitabine/*nab*-paclitaxel had similar weight change trends to those treated with mFOLFIRINOX (Fig. 3). Necropsy analysis revealed that mice treated with gemcitabine/*nab*-paclitaxel died because of disease progression, without significant treatment toxicity. We note that mice treated with two cycles of gemcitabine/*nab*-paclitaxel gained weight between diagnosis and death, which we attribute to abdominal ascites and tumor progression. Additionally, In that regard, it seems that gemcitabine/*nab*-paclitaxel could be safely considered in KPC mice with spontaneous pancreatic tumors.

Despite the improved outcomes in mice treated with gemcitabine/*nab*-paclitaxel, the mice cohorts treated with mFOLFIRINOX still showed reasonable OS (median OS, FFX X1: 11 days, FFX X2: 15 days) (Fig. 1A). Moreover, mice treated with one or two cycles of mFOLFIRINOX had longer OS compared to mice in the control cohort (Fig. 2C). As such, mFOLFIRINOX can also be safely considered in KPC mice. This feasibility study was not designed to assess potential futility of mFFX compared to gemcitabine/abraxane, but we do note that the there was a survival trend in the latter combination with a more favorable toxicity profile, as has been reported in a recent Phase II study [21].

Lastly, we analyzed the use of chemoradiation with SBRT in KPC mice, and we show that the addition of local treatment with radiation therapy to chemotherapy further improved OS (Fig. 2C). This goes in line with pancreatic cancer treatment in humans, where local therapy with surgery and/or radiation therapy has been shown to improve survival and recurrence rates [22, 23]. Furthermore, KPC mice tolerated SBRT treatment well, with only one mouse dying of radiation-induced toxicity (Table 4). SBRT treatment has become increasingly common in clinical practice, and as such, incorporating chemoradiation with SBRT in pre-clinical models would help to best parallel clinical management. In that regard, our data show that mFOLFIRINOX, *nab*/paclitaxel, and SBRT can be safely used and combined in KPC mice.

## Conclusions

Our data show that the use of mFOLFIRINOX and gemcitabine/*nab*-paclitaxel is feasible in spontaneous pancreatic tumors in KPC mouse models. Mice receiving two cycles of chemotherapy had improved survival, and did not show signs of severe toxicity. We further show that chemoradiation using mFOLFIRINOX or gemcitabine/*nab*-paclitaxel chemotherapy regimens with SBRT is also feasible, and was also well tolerated in KPC mice. Our results should encourage future pre-clinical studies to incorporate modern chemotherapy combinations in order to best mimic the current clinical standard of care.

## List of Abbreviations

FFX: FOLFIRINOX
GEM/AB: gemcitabine/*nab*-paclitaxel
IQR: interquartile range
mFFX: modified FOLFIRINOX
OS: overall survival
SAIF: Small Animal Imaging Facility
SBRT: stereotactic body radiation therapy

## Declarations

### Ethical Approval

The University of Texas MD Anderson Cancer Center Institutional Review Board approved all protocols in this study. All mouse work was approved and done in accordance with the Institutional Animal Care and Use Committee of The University of Texas MD Anderson Cancer Center under protocol IACUC #00001252-RN02. The studies were carried out in compliance with ARRIVE guidelines.

### Consent for Publication

Not applicable.

### Availability of Data and Materials

Research data are stored in an institutional repository and will be shared upon reasonable request to the corresponding author.

### Author’s Contributions

AMD, JAJ, and CMT were responsible for the concept and study study design. AMD performed all the mouse experiments and statistical analysis performed by JAJ. All authors contributed substantially to the writing of this manuscript, through performing literature review, data analysis, data interpretation, manuscript drafting, and providing significant comments and/or edits to the final manuscript.

## Acknowledgments

We would like to thank the University of Texas MD Anderson Cancer Center Institute for Applied Cancer Science (IACS) for generously providing the chemotherapy used in our study. This research is supported in part by the National Institutes of Health through MD Anderson’s Cancer Center Support Grant CA016672. This work was also supported by the NIH/NCI under award number P30CA016672 for use of the Small Animal Imaging Facility.

