## Supplementary figures and images for "Feasibility of Administering Human Pancreatic Cancer Chemotherapy in a Spontaneous Pancreatic Cancer Mouse Model"

### Supplemental Figure 1

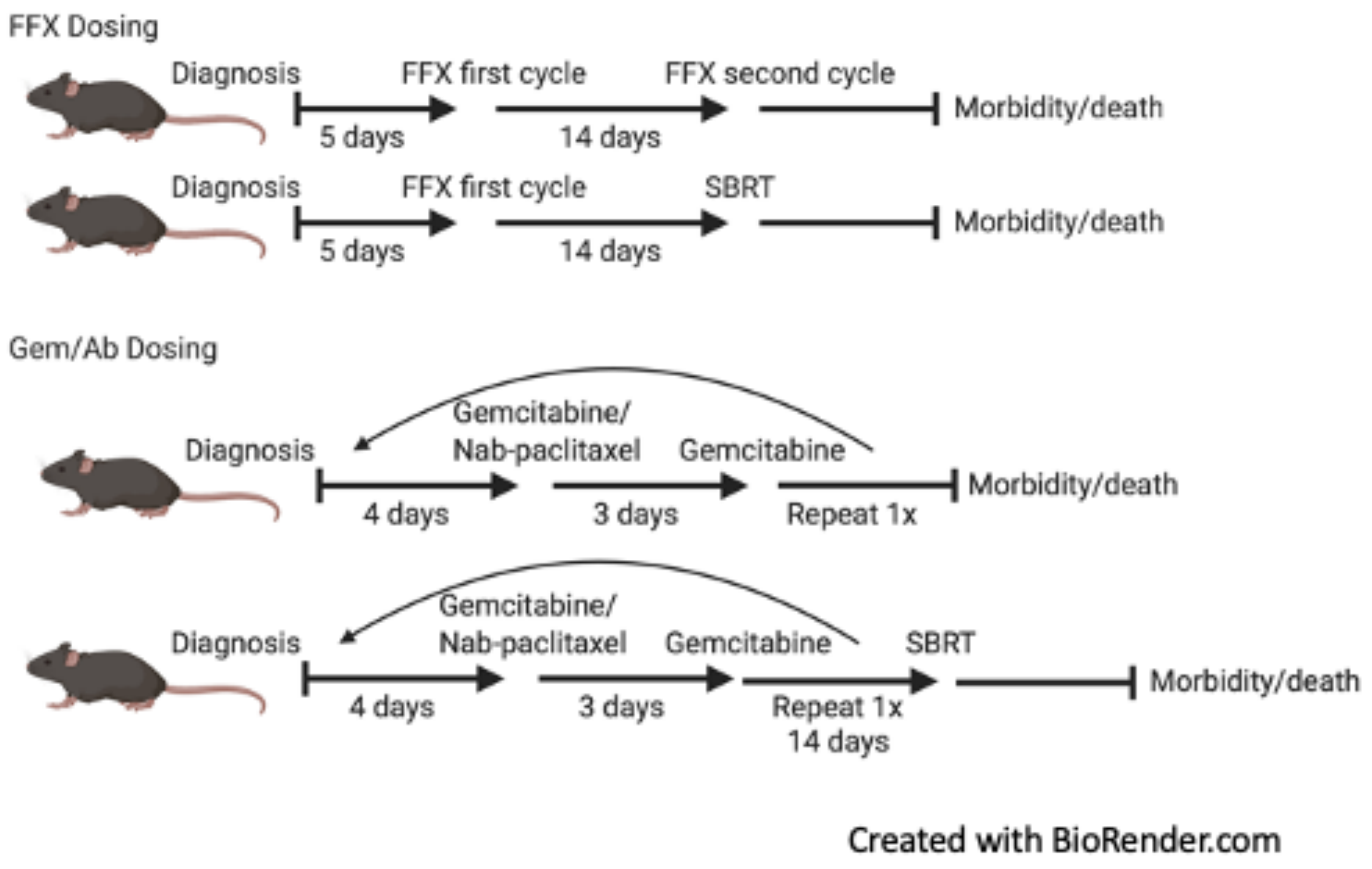
